# Learning for angling: an advanced learning capability for avoidance of angling gear in red sea bream juveniles

**DOI:** 10.1101/2020.01.30.925875

**Authors:** Kohji Takahashi, Reiji Masuda

## Abstract

Angling has been the cause of mortality for fish since ancient. The avoidance learning for angling gear could be considered as a survival strategy against the mortality by angling. Whereas some studies indicated the possibility of avoidance learning for angling gear, most studies investigated the avoidance learning by using groups of fish, in which it is difficult to reveal the process and mechanisms of the learning. The present study elucidated the avoidance learning for angling gear by experiment of single fish in a tank using red sea bream *Pagrus major* juveniles. Individuals with only once or twice of experience for angling avoided angling gear while showing the feeding motivation for pellets, representing avoidance learning for the angling gear. Most of the experienced individuals avoided the krill attached with a fishing line, but not krill and pellets near the angling gear. Feeding rate for prey on a fishing line at two month after the angling trial demonstrated that approximately half of fish kept the memory for angling gear. A series of experiment for angling gear elucidated that red sea bream juveniles are equipped with considerable learning capability for angling gear, suggesting a cognitive evolution for angling.

## 1 Introduction

Angling has been conducted as a major fishing method since ancient [1, 2]. Recent study indicated the evidence that hooks for angling had been made at the 35,000–30,000 years before present [3]. Also, angling is one of popular leisure around the world. A study in Japan showed that one in eleven people enjoy angling as a leisure [4]. It is predicted that angling continues to be central in fishery societies and economies around the world.

On the other hand, the hooking and capturing during angling process is a matter of life and death for fish. Some studies showed the evidence that angling leads to the mortality of fish [5–7]. Even if an angling does not lead to death for fish, the angled fish would suffer from the physiological damage, behavioral change and stress by the angling [8–10]. For example, large mouth bass changed the nest site for parental care by multiple times of angled experiences [11]. Line fishing would accelerate the natural selection for fish which is targeted by fishing [12], and thus they should equip the capability to avoid angling gear efficiently.

The avoidance learning for angling gear is considered as a survival strategy against the angling, because fish should acquire the information of angling gear as a dangerous subject in fishing ground where they frequently encounter angling gears. Past studies suggested that fish can avoid the angling gear through the experience being angled [13–16]. Some studies showed that the angling rate in a natural condition and fish pond is declined over time with angling [17–20]. However, most of the past studies had investigated the avoidance learning for angling by experiment using groups of fish in a large scale, e.g., ponds or large experimental tanks. There are some problems to elucidate the learning of angling gear in such a scale of experiment. For example, the feeding motivation should be confirmed for an individual avoiding the angling gear, because the motivation may be lowered by the stress from angling experience for not only angling gear but also prey itself [21,22]. Also, aversive experience often enhances the alertness of fish [23]. If an individual to be angled enhances the alertness, the fish would be less vulnerable to angling than naive fish, because bold fish often feed more aggressively than fish being cautious [15,24]. Conversely, the competition with conspecific mates often prompts their feeding motivations within a shoal [25,26]. Thus, fish may feed the angling gear in a group even though they had learned the angling gear as a dangerous subject.

In particular, it is unclear for the performance and mechanism of avoidance learning for angling gear, because it is difficult to elucidate these factors by experiments using groups of fish. Animals should recognize appropriately and quickly essential information to survive, and they often equip the cognitive ability fitted to the ecology [27, 28]. Learning performance for angling should be evaluated in consideration of learning process and retention in individual basis, i.e., how many times of angling experience are required to learn the avoidance of angling gear, and also how long it is retained on the memory for angling. Also, a pattern of an associative learning often depends on environmental factor they originate [29–32]. To understand the mechanism of learning for angling, it is important how fish learn the angling gear as an aversive object.

The present study investigated the avoidance learning for angling gear by using red sea bream *Pagrus major* juveniles. The species is one of the most important fishes for fisheries in Japan, and hatchery-reared fish are available from commercial fish farms. The experiment was conducted by a singular fish in experimental tanks to eliminate the effect of group. First, the learning process and the retention of learned information for short period were investigated by presentations of angling gear on consecutive 28 days. Second, we investigated what is the key of avoidance learning by presenting angling gear separately. Third, we compared the angling rate between experienced and control fish, which had never been angled, by blind test. Thereby, we evaluated the effect of the learning for the avoidance of angling gear. Finally, the long term retention of learned information was tested for each individual.

## 2. Material and methods

### (a) Materials

Twenty two hatchery reared red sea bream juveniles were used for experiment (details in electrical supplemental material). Eleven blue polypropylene containers (width 45 × length 66 × depth 33cm, depth 25 cm) were used for the experimental tank (details in electrical supplemental material). A fish was introduced in each tank, and each fish was acclimatized over three days with feeding cycle in the morning (7:00-10:00) and evening (14:00-17:00) everyday. The experiment was initiated when the fish foraged pellets within 30 s after feeding in the morning.

### (b) Experiment 1: the process of learning for angling gear

An experimental trial consisted of a feeding motivation test and a presentation trial. The feeding motivation was confirmed by providing pellets (three to four pellets), and when the fish ate the pellets within 30 s, a presentation trial was immediately conducted. By confirming the feeding of pellets before presenting angling gear for each trial, we evaluated whether the fish would avoid the angling gear while having the motivation of feeding.

Individuals were divided into either of angling treatment and control (11 fish was used for each treatment). Angling treatment fish was presented with the angling gear (details in electrical supplemental information), and the feeding behavior for a krill attached with the angling gear was observed for up to a maximum of 60 s. When fish took a krill with hook into the mouth, the fish was quickly captured out of the water through the angling leader, i.e., the fish was angled. The angled fish was kept in air for at least 30 sec, and then returned to the experiment tank after removing the hook with the snout. When it was difficult to remove the hook from mouth, the leader was cut; i.e., the hook was remained in mouth of the fish, but it did not affected feeding motivation of fish. If fish did not feed a krill for 60 s, the presentation trial was ended as “no angling”. Also, biting times, i.e., the number of time that fish pecked the krill without intake by mouth, were counted during the observation period. When the fish took the krill away without being hooked during the observation, another krill was attached with the angling gear. Control fish was provided with a gear of which hook end was cut by pliers; thus, they could feed a krill attached with the gear without being hooked. An experiment trial for each treatment was conducted during daytime before noon (7:00-11:00). To reduce the variability of hunger level for each individual, sufficient pellets were fed for both angling and control fish every afternoon (15:00-18:00). The experimental trial was repeated for consecutive 28 days. The experiment was replicated for two cycles (1st cycle: angling treatment, n=6, control, n=5; 2nd cycle, angling treatment, n=5, control n=6).

We analyzed the effect of angling treatment for feeding behavior (“fed (including biting without hooking)” or “not”) by generalized linear mixed model (GLMM) with the “lme4” package [33] for R statistical software. The error distributions of the response variables were fitted to the binomial distribution, using restricted maximum likelihood parameter estimation. The fixed factors were “treatment (angling or control)”, “trial (1-28 trials)”, and “interaction (treatment × trial)”, whereas “individual” was treated as a random factor because the feeding behavior of each individual was repeatedly measured during 28 trials. To evaluate the behavioral change of angling treatment, the angling and feeding of only angling treatment fish was fitted to GLMMs (variable factor: angling: “angled” or “not”, feeding behavior: “fed” or “not”, fixed factor for each variable factor: “trial”, random factor: “individual”, error distribution: binomial). Biting time was also fitted to GLMM with the error distribution of Poisson (fixed factor: “trial”). To evaluate the feeding behavior while avoiding angling, angling rate focused on only the fish which showed the feeding behavior for angling gear was also fitted to GLMM (fixed factor: “trial”, error distribution: binomial). We used the Wald test to evaluate the effect of the fixed factors for each model.

The learning performance was investigated for each individual in the angling treatment. When a fish avoided the angling even though it fed on pellets in the motivation test, the fish was considered as leaned. The trials to be required until the learning was investigated for each individual. After the learning, the retention of the learned information was evaluated by days to be re-angled as a short duration memory.

### (c) Experiment 2: the mechanism of learning for angling gear

Each individual was tested to investigate what of the angling gear was avoided by the learned fish after one hour of the Experiment 1. In the test, both of angling and control treatment fish were provided with following angling gears: “pellets and angling gear”, “krill with the line”, and “krill” (electronic supplemental material, figure 2a). The “pellets and angling gear” without krill was presented to evaluate the decrease of the feeding motivation of fish by providing the angling gear. In the “krill with the line” test, an angling gear was presented with a krill attached on a line without a hook: i.e., the same gear as one of control treatment. The “krill” test was the presentation of a krill without angling gear to confirm the avoidance for prey itself. The feeding motivation for pellets was confirmed just before each test. These presentation tests were conducted sequentially for each individual to reduce the number of fish used in the experiment. The order of presentation test was fixed to decrease the effect of previous test. Each presentation test lasted for 60 s or until fish foraged the pellets or krill. The feeding rate for each presentation was compared between treatments and among presentation tests in each treatment by fisher’s exact test. The latency to feed was measured for each test, and compared between treatments by Student’s *t*-test.

### (d) Experiment 3: the effect of learning for angling gear

At one hour after the end of the Experiment 2, the angling test was blindly conducted by third persons who were not informed about the experiment information. The blind tests were conducted by eight participants in total. In the test, the angling gear attached with a hook and krill was presented for 60 sec in the same manner as presentation trial of angling treatment in experiment 1. The feeding behavior was observed during the test, and then the latency until angling was measured if the fish was hooked. The angling rate was compared between angling and control treatment by Fisher’s exact test, and the odds ratio of angling for angling treatment was estimated against control. The fish was then captured after the test, transferred into the beaker, and was photographed to measure the body length using Image J software (Open Source, Public Domain, NIH). We confirmed that there was no difference of body length between treatments (angling treatment, 117.8 ± 8.8mm (mean±standard deviation); control, 118.8 ± 12.6 mm; Student’s *t*-test, t_1, 20_=−0.19 p>0.84). Then, individuals of angling treatment were introduced into rearing tanks (200L transparent circular tank) for each experimental cycle, and they were reared in the same manner as the stock tank until Experiment 4.

### (e) Experiment 4: the long term retention of learning for angling gear

At about two month (87-98 days) after the Experiment 3, the retention of learning for angling gear was investigated for each individual of angling treatment fish (n=10, because a fish of 2nd cycle died before the test). On the day before testing, a fish was introduced in an experimental tank to be acclimatized for a night. After a night, each fish was tested by presentation of “pellets and angling gear” and “krill with the line” in the same manner as in the Experiment 2; the feeding motivation for pellets was confirmed before each presentation. The feeding rate was compared between presentation tests by Fishes’ exact test.

## Results

### (a) Experiment 1: the process of learning for angling gear

All the fish, both angling treatment and control, accepted the presented pellets in the feeding motivation test. Thus, lack of feeding with the presence of angling gear on a presentation trial was considered as avoidance learning for angling gear, but not by loss of feeding motivation.

For the feeding behavior in both treatments, there were significant effects of treatment, trial, and interaction (treatment × trial) in the GLMM analysis (all factors P<0.001; figure 1 & electronic supplemental material, table S1a). For the angling treatment fish, there was a significant effect of trial in each GLMM (feeding behavior P<0.01, angling P<0.001, biting times P<0.05, electronic supplemental material, figure S2,3 & table S1b). Evaluating the angling of fish representing feeding, there was a significant effect of trial for angling rate (table 1c, P<0.001; electronic supplemental material, figure S4 & table S1c).

**Fig 1.**
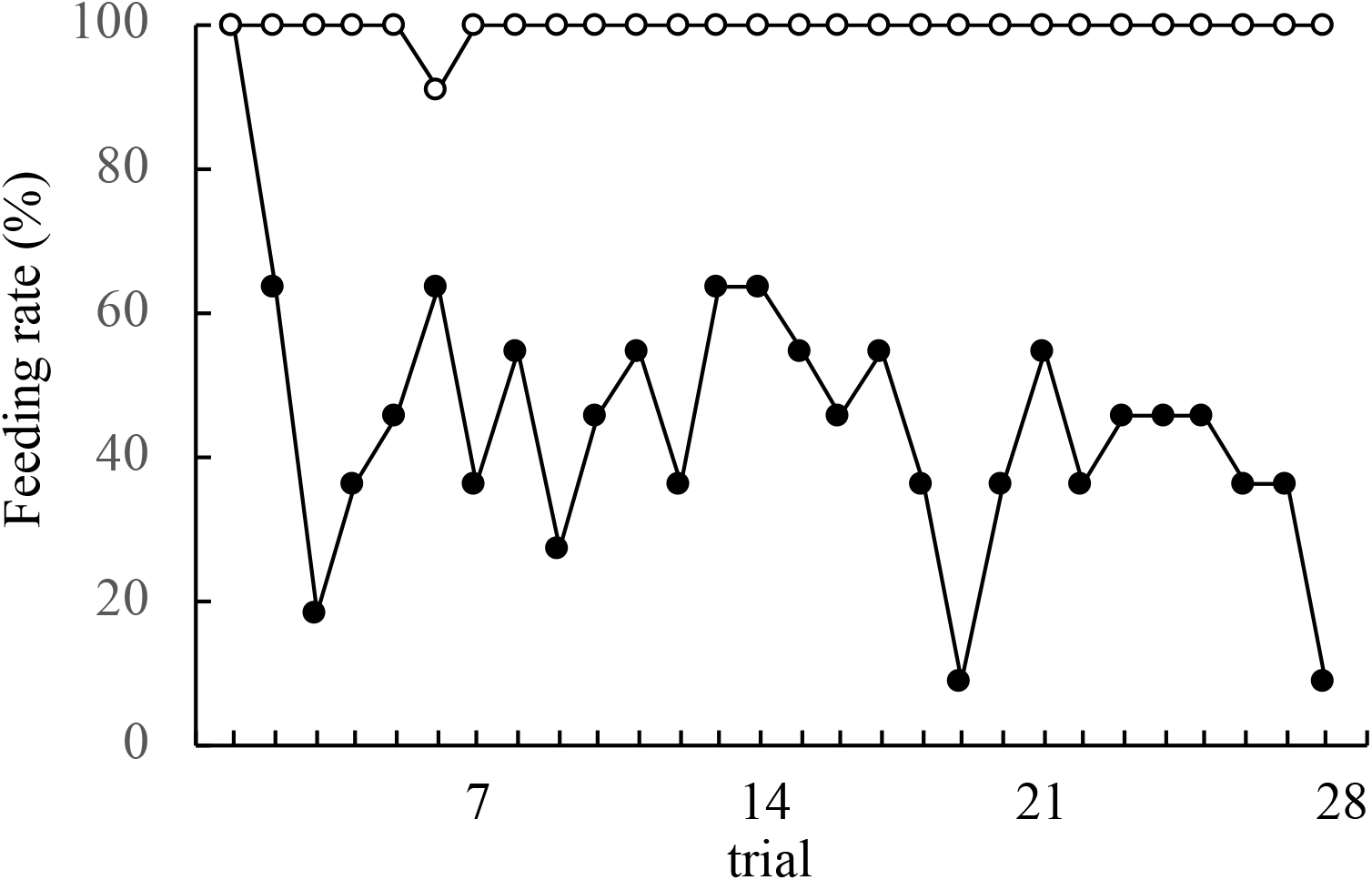
The feeding rate of fish in the control (open circle) and angling (closed circle) treatment during the 28 trials in Experiment 1

For individual analysis of angling treatment, all individuals avoided the angling gear after one (n=5) or two angling trials (n=6); i.e., fish required 1.6 ± 0.5 (mean ± standard deviation) trial to learn the avoidance of angling gear. Meanwhile, all fish was re-angled within 13 days after the avoidance learning (electronic supplemental material, fig s5), and then 4.3 ± 3.3 days were required until next angling. Times of angling during 28 days varied from three to eight among individuals.

### (b) Experiment 2: the mechanism of avoidance learning for angling gear

Angling treatment fish showed the feeding rate of 100 % (11/11 individuals) for “pellets and angling gear”, 9.1 % (1/11) for “krill with the line”, and 100 % (11/11) for “krill”, respectively (figure 2b). The feeding rate of control fish was 100 % (11/11) for all the presentation test. The feeding rate of “krill with the line” for angling treatment was significantly lowered than control treatments (P<0.001), but not different from others (P=1.00). For angling treatment fish, the feeding rate of “krill with the line” was significantly lower than other presentation (P<0.001). There was no difference of feeding rate among presentation tests in control (P=1.00). The latency to feed “krill with the line” of control was faster than the fish (11 sec) of angling treatment (one-sample *t*-test, t_1, 10_=−19.53, P<0.001; electronic supplemental material, fig s6). Meanwhile, there was no difference of the feeding latency between treatments for “pellets and angling gear” and “krill”(“pellets and angling gear”: t_1, 20_=−1.33, P<0.20, “krill”: t_1, 20_=−1.04, P<0.31).

**Fig 2.**
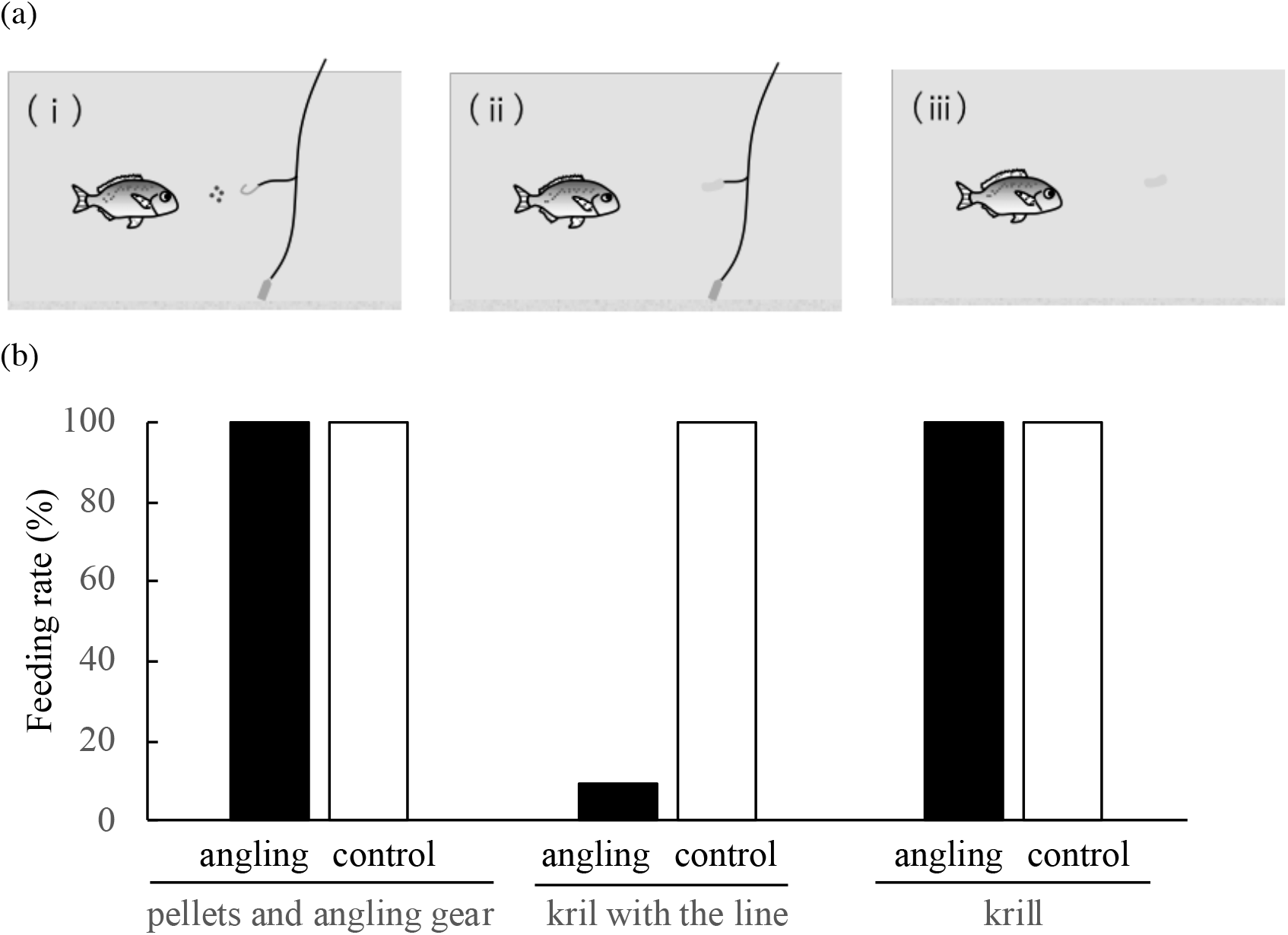
(a) Schematic illustrations of test presentation in experiment 2: (i) pellets and angling gear, (ii) krill with the line, and (iii) krill. (b) The feeding rate of fish in the angling (black column) and control (white column) treatment when three types of gears or prey were presented in Experiment 2

### (c) Experiment 3: the effect of learning for angling gear

The angling rate in blind test was 18.2% (2/11) for angling treatment and 81.82 % (9/11) for control, respectively, and there was a significant difference of angling rate between the treatments (P<0.01). The latency for angling was 18s and 52s for angling treatment, and 9.4 ± 10.2s for control, respectively, and there was a significant difference between treatments (one-sample *t*-test “control vs 18s”; t_1, 8_=−2.37, P<0.05). The cause of not being captured in angling treatment was either they did not feed at all (in two individuals) or showed only biting (in other seven individuals). For control fish, two fish showed the biting for the angling gear but were not captured. The odds ratio for angled fish in the angling treatment was 0.05 against control.

### (d) Experiment 4: long term retention of learning for angling gear

The feeding rate was 100 % (10/10) with the presentation of “pellets and angling gear” and 50% (5/10) with that of “krill with the line”. The foraging rate of “krill with the line” was significantly lower than “pellets and angling gear” on the two month later test (P<0.04).

## Discussion

All individuals including both treatments of angling and control fed the krill attached with angling gear and were angled on the first trial. Meanwhile, angling treatment fish decreased to feed from the angling gear with the experience of angling, although they had always demonstrated the active feeding on the pellets in the motivation tests prior to the angling trial. This result suggests that the fish recognized the angling gear as an object to avoid feeding through the experience of angling; i.e., avoidance learning for angling gear. Whereas past studies using groups of fish suggested the avoidance learning for angling gears [13–16], these studies have not confirmed the feeding motivation of fish during experiment. The present study verified the avoidance learning for angling gear to elucidate the fact that fish desired prey at the presentation of angling gear.

The angling rate in the angling treatment decreased during 28 days, suggesting that learning progressed gradually as the experience of angling. The major factor for the decrease of angling would be the refusing to feed from the angling gear, because the feeding rate also decrease with trials during angling treatment. Meanwhile, the feeding for angling gear was maintained to a certain extent even after the learning progressed. Or rather, biting times, i.e., pecking krill attached with the angling gear without being hooked, had increased during learning process. This means that fish getting experience of angling might become not only to avoid angling gear, but also to steel prey on the angling gear without being hooked. In fact, the angling rate regarding only feeding fish decreased remarkably with trials. The present study indicates that red sea bream juveniles improve the feeding behavior of angling gear while avoiding to be angled. Improvement of feeding skill is found in fishes [34, 35]; e.g., the shooting accuracy in archer fish was improved rapidly through the experience of target movement [36]. However, in our knowledge, present study is the first to have elucidated the improvement of feeding for angling gear.

All individuals of angling treatment learned to avoid angling gear through only once or twice experience of angling. Red sea bream juveniles were able to quickly learn the angling gear as a dangerous object. Similarly, past studies suggested that the fish can learn angling gear by only a few times of angling experience [14,15]. For example, Beukma [37] found in carp by an experiment on artificial lake that many individuals became invulnerable to angling once being angled; however, this study could not rule out the possibility of reduced feeding motivation. Avoidance learning for angling gear may be a simple task for fish, at least for some species which are subject to angling. Essential information in an ecology is often learned rapidly with only a few experience in fishes [38, 39]. For example, predator information is learned by a single conditioning [40]. Shanny, which form the nest in rock tide pool, can learn the location of shelter with only one trial [41]. The remarkable cognitive ability for angling gear in red sea bream implies that the learning for angling gear is important in their ecology.

Although angling treatment fish quickly learned to avoid angling gear, all individuals were re-captured after the learning; 4.3 days on average were required to be re-angled. This means that by no means fish can avoid completely the angling gear after the learning of angling. Fish may be angled in a place where anglers are abundant, such as a major fishing ground or fishing pond, even if the fish can quickly learn the angling gear as dangerous subject. Some studies have shown the re-angling of fish that had previously experienced angling [42, 43]. For example, Tsuboi & Morita [44] using white-spotted charr found that some of fish were angled repeatedly during research period. The study also showed that the fish with a greater experience of being angled were more angled; that is, there were individual variations of learning for angling. In the present study, angling times varied among individuals of angling treatment, and some fish were vulnerable to angling. In the following experiment of the present study, the vulnerability of angling is related to the boldness of individuals (Takahashi, unpublished); i.e., bold fish were angled more frequently than shy ones. The individual difference of learning should be investigated in more detail. On the other hand in the present study, the fish is confined in a small tank, and then they were obliged to be encountered with angling gear after the learning. Thus, they might have fed the prey on the angling gear knowing that it was dangerous; the increased biting times of angling gear during angling trials supports this speculation.

Although angling treatment fish was re-angled in a few days after the fish had once learned angling (Experiment 1), the fish developed the cognition for angling by repeated presentation of angling gear. At two months after the 28 consecutive angling presentations, approximately a half of angling treatment fish avoided a krill attached with a line (experiment 4). This suggests that the angling treatment fish, at least a part of them, retained the learning for angling gear even if they had not encountered the angling gear for this period. Similar suggestions of long term memory for angling are also found in experiment which was conducted in a large pond [37]; the experiment suggests that fish kept the memory of being hooked for at least a year. The present study has verified the long term memory of avoidance learning for angling gear by a small scale experiment. Retention of learned information can be often related to the ecology of animal. For example, food-storing birds have better spatial memory than birds that do not store food [45]. In fish, the retention of information for predator is shaped by individual growth rate as a behavioral tactics [46]. For memory retention of red sea bream, Kaneko [47] showed that most fish were not able to retain the learned information of feeding area for 60 days. It might be important for red sea bream to keep the information of angling gear for a long term, because they would often encounter the angling gear in their lives after the learning for angling.

In Experiment 2, all of learned fish fed pellets under the presence of angling gear. It is expected that the feeding motivation was not lowered by the vigilance against the presence of angling gear. Also, all fish fed a krill without the line and hook. Some study found that fish avoid prey containing aversive substance, such as toxic or unpalatable food [48,49]. However, the fish in the present study did not recognize the krill itself as a prey which should be avoided. The result indicates that the avoidance learning for the angling gear is regarded as distinct from food aversion learning. Aversion learning for prey is often affected by combination between conditioned stimulus and unconditioned stimulus; for a classical example of rat, the aversion for prey can be conditioned with lithium chloride, but not with electrical shock [50]. The aversion learning for angling in the present study would be difficult to be associated with prey itself in red sea bream.

Meanwhile, almost all fish of the angling treatment avoided to feed a krill with a line; the feeding latency of a feeding fish was longer than control feeding fish. This means that red sea bream juveniles evaluated the risk of angling gear with the presence of fishing line attached with a prey. Whereas past studies suggested that fish can avoid angling gears through a learning effect, the mechanisms have not been clear [16]. The experiment elucidated that a prey attached with a fishing line is the key factor of the learning for angling gear. The result in experiment 2 indicates that they could feed a prey they had once eaten in angling process unless the prey is attached with a fishing line. Angled fish are often returned to fishing ground, either being released intentionally by angles or the fish being able to escape during the angling process. The fish after the release must avoid a prey which lead to be angled, because repeated angling would increase the risk of mortality [5–7]. However, while avoiding the dangerous prey, the fish need to take prey for their lives. If fish learn to avoid the prey itself previously angled, such as food aversion learning, the fish would lose a chance to get the prey even when the prey is safe without angling line. It would be essential to take a prey discriminating correctly whether the prey is dangerous. The making decision of feeding a prey, i.e., estimating a presence of angling line, must be useful for fish to survive in a fishing ground.

In the blind test after the angling treatment (experiment 3), the angling rate of angling treatment fish was remarkably lower than control fish; twenty times difference of the vulnerability for angling between treatments. The experiment showed that the vulnerability for angling is markedly improved by repeated angling experiences. There are some possibilities for the decline of angling in large scale experiment. For example, the recapture rate of angled fish decreases if fish leave the fishery ground [51]. Stålhammar [22] found that the feeding behavior for a prey of pike was delayed by an angling experience. Meanwhile, angling treatment fish in the present study were not able to go away from the angling gear in the experiment tank, and had the active feeding motivation for prey just before the angling test. Thus, it is predicted that the angling vulnerability was improved by the avoidance learning for angling gear. Re-angling of learned fish should have occurred less in a natural fishing ground than in a limited space on the present study. Furthermore, the learning efficiency is often enhanced by the social learning for various contents [52,53]. Whereas it is not still clear for the social learning of angling gear, if the social learning for angling gear is established in groups of fish, the invulnerability of angling would be prompt in natural condition.

Animals often have a cognitive ability adapted to their ecology [27,28], and the adaptive cognition is often prepared innately in them [29]. For example, tropical poeciliid in higher predation pressure exhibit an innate behavioral character adapted with predatory environment [54]. Also, hatchery-reared jack mackerel juveniles under rearing condition develops ontogenetically the learning capability to fit with the habitat shift during life history in nature [55]. The red sea bream juveniles in the present study were hatchery-reared, and thus had never encountered angling gear in their lives before the experiment. Nevertheless, they demonstrated the advanced learning capability for angling gear. This suggests that the cognition for angling gear is innately equipped in the red sea bream juveniles. Red sea bream at the adult stage is targeted for fishery of pole-and-line fishing and recreational angling in Japan [56]. The juveniles are released for stock enhancement around Japan to coastal zone and frequently captured by recreational angling, but forced to release for regulation of angling size by fishery adjustment rule in each prefecture [57] (see also Kanagawa prefecture Web page, http://www.pref.kanagawa.jp/index.html). It is predicted that they repeatedly encounter angling gears during the life. The advanced cognition for angling would help them to survive after the release.

In Experiment 2, the juveniles learned angling gear depending on a line attached with a prey. It is considered that such learning would not be expected in their lives except for angling process, because a prey with a line would not be present in the natural environment without angling gear. The avoidance learning in red sea bream might be formed under a cognitive mechanism specialized in angling. Angling has been a common fishing method since ancient times [1–3], and then fish had been exposed to the risk of angling on their evolution. The cognition for angling gear might be evolved in red sea bream under the selection for angling. Fishing has the potential to induce evolutionary change in traits in fish populations [12, 58, 59]. For example, faster-growing genotypes which may be more vulnerable to fishing depletion were more frequently harvested than slow-growing fish [60]. In largemouth bass, recreational angling induced evolutionary changes in various physiological and behavioral traits after only four generations [61]. The adaptive cognition for angling gear in red sea bream might have occurred through the natural selection by the angling. The present study suggests the cognitive evolution for a fishing activity. Whereas the avoidance learning for angling gear is suggested for various fish species, past studies are limited to the experiment in large scale [13–16]. These studies focused on only the targeted species for angling, including the present study. It is unclear for the learning capability of angling in non-targeted species. In future, comparative study between targeted and non-targeted species would elucidate the cognitive evolution for angling in fishes.

## Acknowledgement

We thank also students of Maizuru Fisheries Research Station (MFRS) which assisted the experiment 3.

## Author contribution

Experiment and writing paper was conducted by Kohji Takahashi, and the experiment was directed by Reiji Masuda.

## Fund statement

The present study was funded by a grant-in-aid for Japan Society for the Promotion of Science Fellows and Early- Career Scientists (KAKENHI 18K14512).

## Ethical notes

All experiments were performed according to the regulations on animal experimentation of Kyoto University. A minimum number of fish was used to test the hypothesis. After the experiment, the fish were donated to local aquarium.

## Data accessibility

Data have been submitted as electronic supplementary material.

