## Supplemental information for "Learning for angling: an advanced learning capability for avoidance of angling gear in red sea bream juveniles"

(a) Fish

Fertilized *P. major* eggs (approximate 20000 eggs), purchased from Marua Suisan Co., Ltd. (Ehime prefecture, Japan), were transported to the Maizuru Fisheries Research Station, Kyoto University, and kept in 500-L transparent polyethylene tanks under the natural light condition. Water in the tank was maintained by exchanging at a rate of 4 L min-1, and an aeration was set in the tank. Sea water used for the rearing tank was pumped up from off the Research Station and fine-filtered. The water temperature was maintained at 20°C using a heater and thermostat. After hatching on May 17, 2013, larvae were provided with rotifer *Brachionus plicatilis* sp. complex (approximately 5 ind/ml), brine shrimp *Artemia* sp. nauplii (approximately 2 ind/ml), and dry pellets (Otohime B1, Otohime C2, and Otohime S2, Marubeni Nisshin Feed Co., Ltd.) with the amount corresponding to growth. Fish were reared for at least nine month in the stock tank, and some of them were used for the present study.

Angling gear

A device of terminal tackle ("Douzuki sikake" in Japanese) was used as an angling gear in the present study; that is, a single hook with a line (approximately 5 cm) was tightened with a fishing leader, and a fishing sinker was attached with the end of the leader (see details in electronic supplemental material, fig s1). The other end of the leader was tied with the tip of a rod (length approximately 90 cm). The barb of hook was pressed by pliers to reduce the stress for hooking and release. The hook was exchanged with a new one when the hook was broken or rusted. A defrosted krill was used as a prey of angling gear. The krill was cut into pieces of approximately 5-8 mm, and then put on the hook.

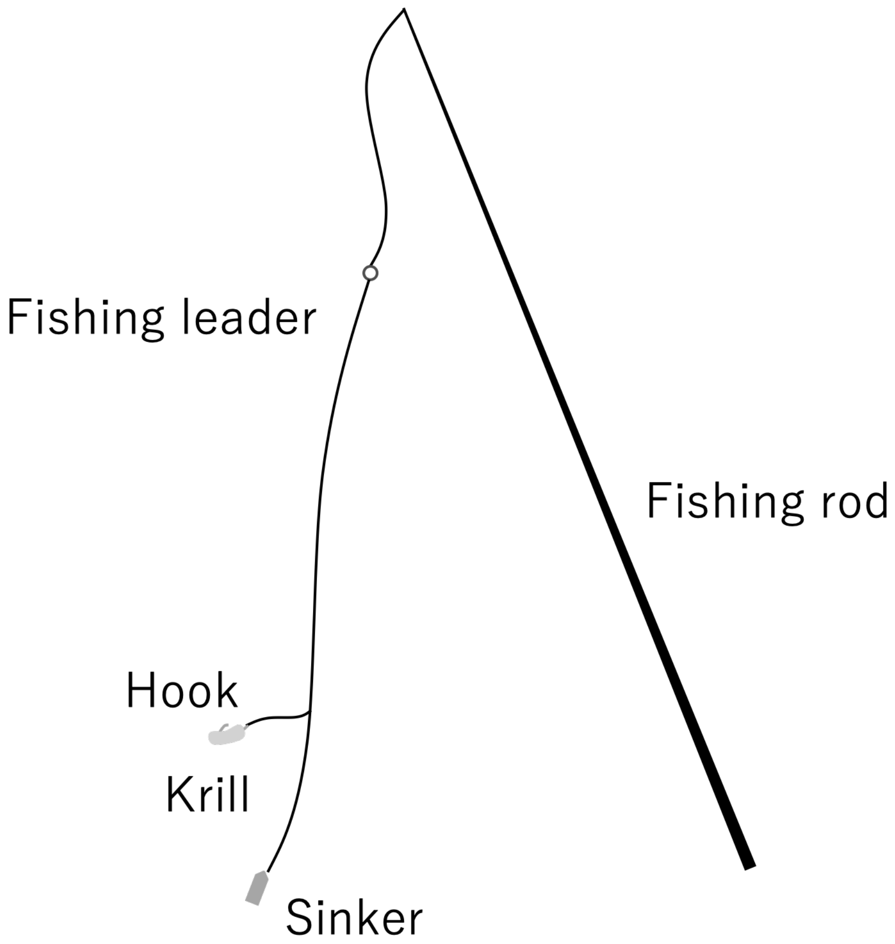


Fig S1 A schematic illustration of angling gear

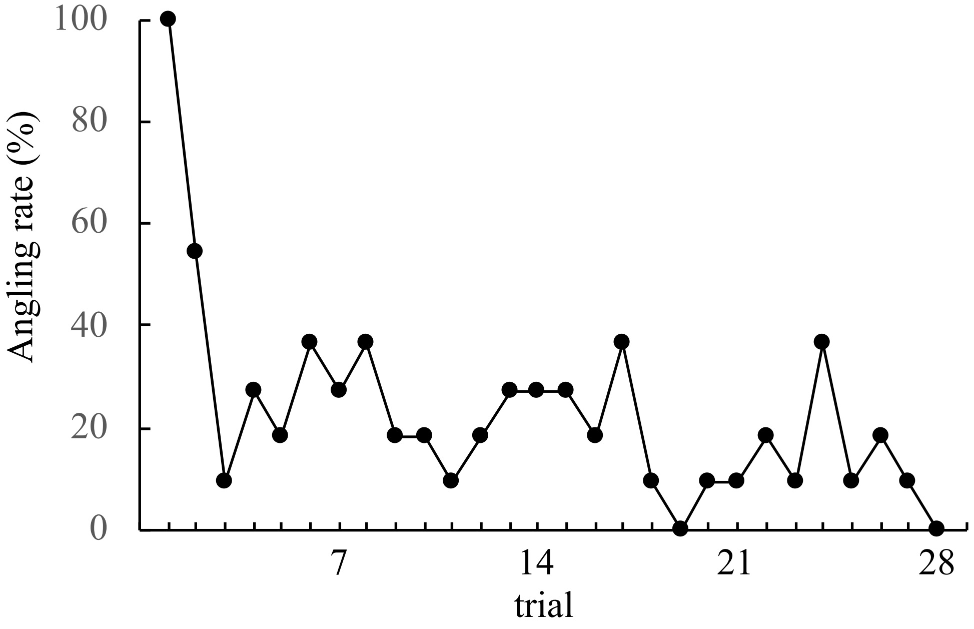


Fig S2 The angling rate of angling treatment fish during 28 trials on experiment 1

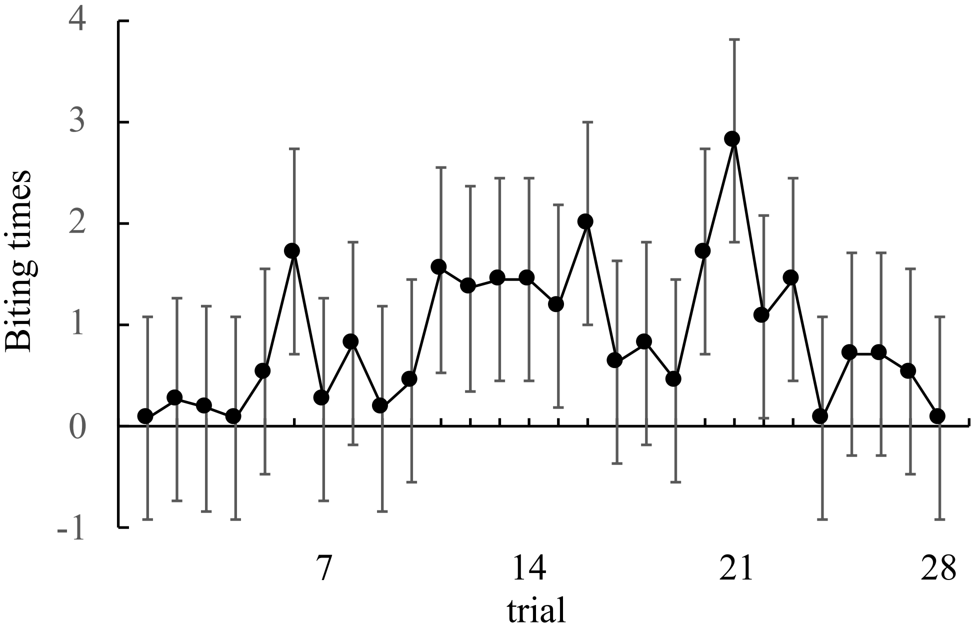


Fig S3 The biting times of angling treatment fish during 28 trials on experiment 1

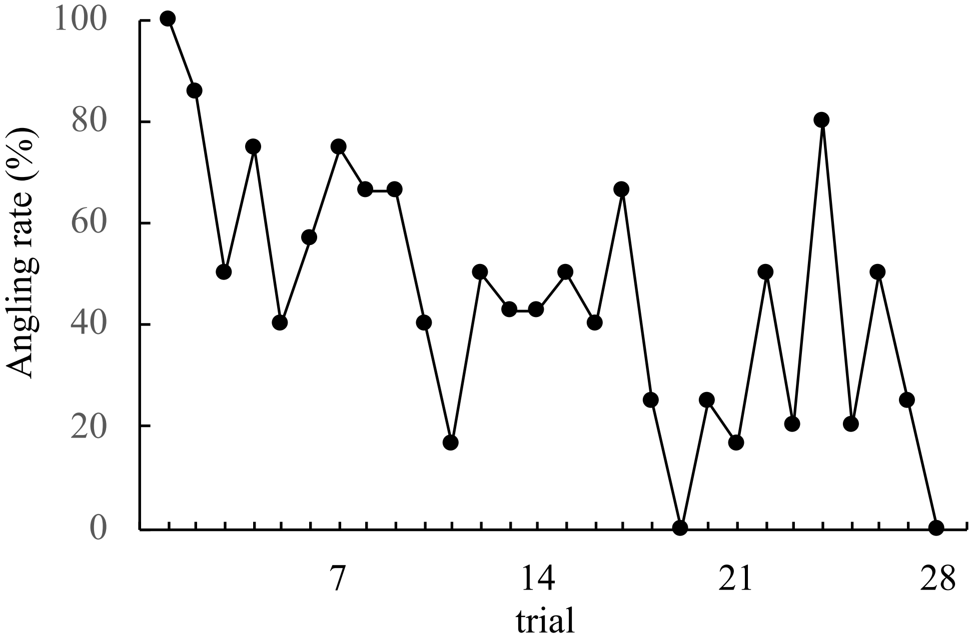


Fig S4 The angling rate of angling treatment fish which represented feeding behavior during 28 trials on experiment 1

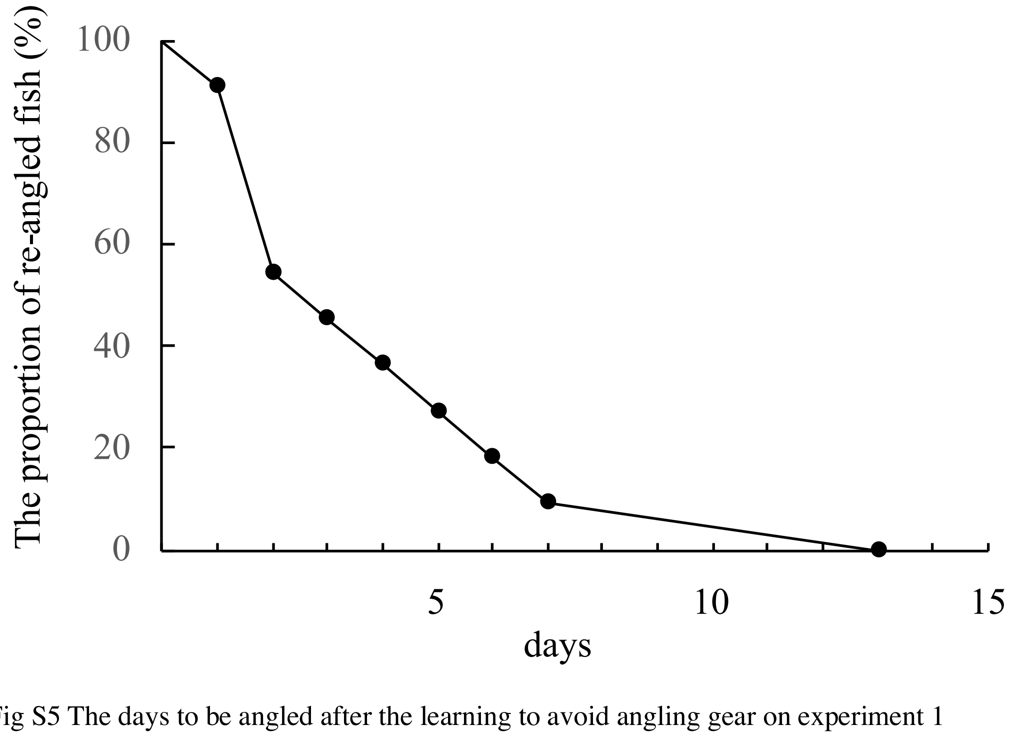


Fig S5 The days to be angled after the learning to avoid angling gear on experiment 1

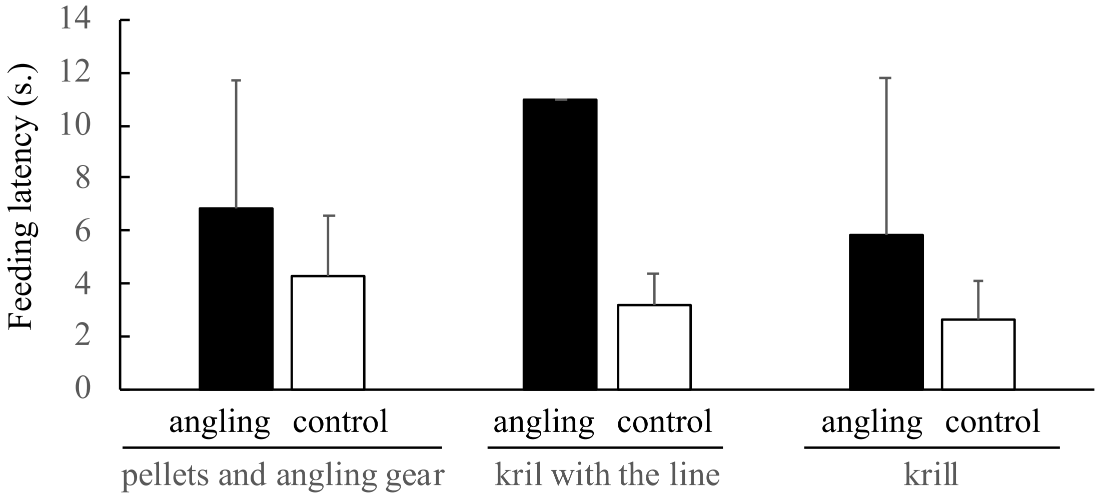


Fig S6 The feeding latency for each presentation test of fish in each treatment on experiment 2. Bars indicate the standard deviation for each data
